# Serial coupling of chromatographic columns having orthogonal selectivity to improve the coverage of oxidised lipidome for mass spectrometry analysis

**DOI:** 10.1101/741579

**Authors:** Alpesh Thakker, Andrew Pitt, Corinne Spickett

## Abstract

Phospholipid oxidation (OxPL) generates a wide variety of products with potentially novel biological activities that may be associated with disease pathogenesis. To understand their role in disease requires precise information about their abundance in biological samples. Liquid chromatography-mass spectrometry (LCMS) is a sensitive technique that can provide detailed information about the oxidative lipidome, but challenges remain. Furthermore, variation in charge of the polar head groups and the extreme diversity of oxidised species make analysis of several classes of OxPLs within one analytical run challenging.

The work in this study aims to develop improved methods for detection of OxPLs by improvement of chromatographic separation through the serial coupling of polystyrene-divinylbenzene based monolithic, and mixed-mode hydrophilic interaction (HILIC) with use of semi-targeted mass spectrometry approaches. The results suggests that by serially coupling two columns, HILIC and monolith, provided the better coverage of OxPL species in a single analytical run. We tested in-vitro generated oxidized species for phosphatidylcholine (PC) and phosphatidylethanolamine (PE) class and the combination of orthogonal chromatographic separation allowed separation of oxdised species from both the classes, which otherwise coeluted.

## 2 Introduction

Global oxidised lipidome analysis of biological samples offers great potential for detection of biomarkers of disease. A single species belonging to the glycerophospholipid group bearing an unsaturated fatty acyl chain can give rise to over 50 different modified species. The species formed through oxidation can further be categorised based on the nature of oxidant, extent of oxidation and concentration of oxidant and other aiding compounds (Reis and Spickett 2012). In lieu of the extreme diversity of phospholipid species due to the fact that more than 1000 of species that can be formed by combination of different headgroups and fatty acyl chains, the number of oxidised species that are generated in response to reactive oxygen species can be overwhelming. In addition, they can exist as many different derivatives and isomers in vivo. This can lead to formation of hundreds of different modified species from the lipidome of mammalian membrane, adding to the complexity and difficulty in characterisation and quantification of these species in biological samples (Nakanishi, Iida et al. 2009). These oxidised phospholipid species (OxPL) can have non-identical biological activities (Bochkov, Oskolkova et al. 2010, Aldrovandi and O’Donnell 2013, Spickett and Pitt 2015). To understand the biological importance of oxidised phospholipids (OxPL) along with their role as a disease biomarker, information on the precise concentration of all oxidised species in biological samples must be obtained (Gruber, Bicker et al. 2012). Moreover, it is important to measure simultaneously all long chain oxidation products including hydroperoxides and hydroxides, and short chain oxidation products including saturated and unsaturated aldehydes and di-carboxylic acids derivatives. This is required to investigate the dynamic assessment of oxidised phospholipids in a disease state, and to examine their relevance to function and disease *in-vivo* (Uchikata, Matsubara et al. 2012).

The analytical technologies to measure the abundance and identification of oxPL species have evolved over the years. In the pre-2000 era, methods involving mass spectrometry measurements were limited and general methodology to analyse oxPL species were used such as UV spectrophotometric assays & fluorescence based assays like FOX2 assay, isoluminol assay and DNPH assay that provided only global measure of oxidative stress without giving any information on the molecular level (Zhang, Cazers et al. 1995, Konishi, Iwasa et al. 2006). Although, these methods provide useful initial insights about potential oxidative state, they suffer from inherent problems related to sensitivity and specificity of detecting individual molecular species, when applied to situations *in-vivo* (Tyurina, Tyurin et al. 2009).

On the other hand, mass spectrometry coupled with liquid chromatography as a technique to measure oxidised phospholipids has gain popularity because it measures mass –to charge ratio (m/z) of compounds and can distinguish or separate different molecular species based on their masses and functional chemistry, therefore, can selectively identify several different species in complex mixture simultaneously. The last decade has seen surge in more sophisticated methods involving use of liquid chromatography coupled with mass spectrometry (LC-MS), supporting the significance of OxPL in health and disease (Spickett 2001, Sparvero, Amoscato et al. 2010, Spickett, Wiswedel et al. 2010, Spickett, Reis et al. 2011, Stutts, Menger et al. 2013).

Due to extreme chemical diversity of the oxidised lipidome together with the range of concentrations of these species in biological samples, complementary analytical approaches are required to monitor it completely (Domingues, Reis et al. 2008, Spickett and Pitt 2015). It is analytically impractical to measure oxidised species of all classes in a single run because of variation in the charge of the polar head group and extreme diversity of oxidised phospholipid species OxPL). Moreover, sensitivity is the major concern, because currently no method is available to selectively enrich or amplify these species to improve their detection (Mousavi, Bojko et al. 2015, Ulmer 2015). Therefore, the strength of the mass spectrometric analysis can be exploited by enhancing the selectivity of the methods.

While mass spectrometer instruments can separate and identify species that have different molecular masses, there are limitations in separation of isomeric and isobaric species (Sandra, Pereira Ados et al. 2010). Moreover, isobaric and isomeric species are not uncommon in oxidative Lipidomics and therefore, to handle the diversity of OxPL species that can be formed, it is often required to couple chromatographic separation with mass spectrometry detection (van Meer 2005, Peterson and Cummings 2006, Wörmer, Lipp et al. 2013).

Various reports describe the combination of either reverse phased and or normal-phase LC coupled with mass spectrometry in multi-dimensional set ups to detect and measure long chain oxidation products(Hui, Chiba et al. 2010, Morgan, Hammond et al. 2010, Thomas, Morgan et al. 2010, Strassburg, Huijbrechts et al. 2012). Most of the methods developed so far relied on mass spectrometric and chromatographic development (Fu, Xu et al. 2014, Groessl, Graf et al. 2015) or focused on high throughput analysis (fast HPLC) at the expense of detection of low abundant oxidation products (Gruber, Bicker et al. 2012). Most of the current methods published so far, have been either confined to a small number of molecular species based on Multiple Reaction monitoring (MRM) based approach or have used high resolution mass spectrometry coupled with liquid chromatography to detect specified oxidised phospholipid species, solely based on accurate masses and/or tandem mass spectrometry (MS-MS) to obtain structural information(Adachi, Asano et al. 2006, Uchikata, Matsubara et al. 2012, Uchikata, Matsubara et al. 2012). The MRM and high resolution based methods are highly sensitive and focused methods and are ideal for observing and quantifying certain pre-determined species. Nevertheless, further improved methods that are capable of quantifying multiple oxidised species in biological samples belonging to different phospholipid class are still required.

To date, there is no report of systematic evaluation of chromatographic separation for OxPL species. The resolution parameter that dictates the chromatographic separation is dependent on three other parameters: efficiency, selectivity, and capacity; the maximum improvement in resolution is related to altering the selectivity parameter, which is the factor that is dependent upon the chemistry of the analyte, mobile and stationary phases (Zhou, Song et al. 2005, Weng 2014). All of these factors may be altered to optimise the chromatographic separation. In this work, one of the primary objectives was to perform systematic evaluation of chromatographic separation by testing several stationary and mobile phase systems to achieve best separation of OxPL species. We also investigated the two dimensional separation by serially coupling two columns that provides orthogonal selectivity. The set up was adapted from (Haggarty, Oppermann et al. 2015) by coupling the HILIC column and reverse phase column.

## 3 Materials and Methods

### 3.1 Chemicals and reagents for liquid chromatography – mass spectrometry (LC-MS)

#### 3.1.1 Chemicals

L-α phosphocholine mixture (PC mixture from egg yolk), L-α phosphatidylethanolamine mixture (PE mixture from egg yolk) and 1,2-diacyl-sn-glycero-3-phospho-L-serine mixture (PS mixture from bovine brain) was bought from Sigma-Aldrich, Dorset, UK. 2-2’-Azobis (2-methylproprionamidine).2HCl (AAPH), a water soluble radical initiator and Sodium Hypochlorite solution (10 – 15 % chlorine) were obtained from Sigma-Aldrich, Dorset, UK. 1-stearoyl-2-oleoyl-sn-glycero-3-phosphocholine (SOPC), 1-stearoyl-2-linoleyl-sn-glycero-3-phosphocholine (SLPC) and 1-stearoyl-2-arachidonyl-sn-glycero-3-phosphocholine (SAPC) were procured from Avanti Polar Lipids, Inc, USA. 1-palmitoyl-2-arachidonyl-sn-glycero-3-phosphoethanolamine (PAPE), 1-O-1’-(Z)-octadecenyl-2-(5Z, 8Z, 11Z, 14Z-eicosatetraenoyl)-sn-glycero-3-phosphoethanolamine (O-18:0, 20:4 PE) and 1-palmitoyl-2-arachidonyl-sn-glycero-3-phosphoserine (PAPS) were procured from Avanti Polar Lipids, Inc, USA. 1-didecanoyl-2-tridecanoyl-sn-glycero-3-phosphoethanolamine(PE (12:0,13:0)), 1-didecanoyl-2-tridecanoyl-sn-glycero-3-phosphoserine (PS (12:0,13:0)) and 1,2-ditridecanoyl-sn-glycero-3-phosphocholine (PC(13:0,13:0)) used as internal standards and were procured from Avanti Polar Lipids, USA. 1-hexadecanoyl-2-azelaoyl-sn-glycero-3-phosphocholine (PAzPC), 1-hexadecanoyl-2-(5’-oxo-valeroyl)-sn-glycero-3-phosphocholine (POVPC) and 1-hexadecanoyl-2-(9’oxononanoyl)-sn-glycero-3-phosphocholine (PONPC) were procured from Avanti Polar Lipids, USA.

#### 3.1.2 Reagents

Deuterated solvents for nuclear magnetic resonance spectroscopy (NMR), namely pyridine-d5, deuterated chloride, deuterated-methanol, and chloroform, were procured from Goss Scientific, UK. All solvents (methanol, chloroform, water, tetrahydrofuran and hexane) were of HPLC grade and obtained from Thermo Fisher Scientific, UK.

### 3.2 HPLC columns

Pro-swift RP-4H (1×250mm) monolithic column was purchased from Thermo Scientific, UK for LC-MS analysis. The Luna C8 column (150mm X 1 mm), Luna C-18 column (150 × 1 mm) and C-30 column (150 × 2.1 mm) were purchased from Phenomenex, UK. Mix – mode HILIC column (300 × 2.1 mm) was procured from HiChrom, UK

#### 3.2.1.1 Optimisation of chromatographic separation using several reverse phase columns and eluent systems

Conventional reverse phase columns like C-8 Luna column (150 × 1mm), C-18 Luna column (150 x 1mm), C-30 Luna column (150 × 2.1mm) and Proswift –RP 4H polystyrene – divinyl benzene coated monolith column (1 x 250mm); and mix mode Hichrom amino based HILIC column (150 × 3.1mm) were evaluated using combination of different eluent systems, while maintaining the linear flow rate and using mass spectrometry for detection. The solvent systems that were used to achieve best separation were: **Solvent system A** consisting of LC-MS grade water with 5 mM ammonium formate and 0.1 % formic acid as eluent A and Methanol with 5 mM ammonium formate and 0.1 % formic acid as eluent B; **Solvent system B** consisting of ternary mixture of methanol, hexane and 0.1 M ammonium acetate (71:5:7) as eluent A and methanol, hexane mixture (95:5) as eluent B; **Solvent system C** consisting of ternary mixture of tetrahydrofuran (THF), methanol and 10 mM ammonium acetate (30:20:50) as eluent A and THF, methanol and 10mM ammonium acetate (70:20:10) as eluent B; **Solvent system D** consisting of 20 % isopropyl alcohol (IPA) in Acetonitrile (ACN) as eluent A and 20 % IPA in 20 mM ammonium formate as eluent B. Gradient run for solvent system A consisted of a 50 minute run with 0 – 4 minute isocratic hold at 70 % B, followed by 3 step gradient increase to 80 % B at 8 minutes, 90 % B at 15 and 100 % B at 20 minutes. The gradient remained constant until 38 minutes and decreased back to 70 % B until 50 minutes. Gradient run for solvent system B was a 45 minute run with 0 – 4 minute at 100 % A followed by 3 step gradient increase to 10 % B at 8 minutes, 40 % B at 20 minutes and 100 % B at 26 minutes with isocratic hold at 100 B until 36 minutes and equilibration to 100 % A from 38 – 45 minutes. Gradient run for solvent system C consisted of a 50 minute run starting at 20 % B at 4 minutes and increasing to 100 % B at 20 minutes, followed by isocratic hold at 100 % B until 36 minutes and back to starting condition at 38 minutes. Gradient program for solvent D consisted of a 45 minute run under HILIC conditions only used for the silica based HILIC column from Hichrom, UK with multi step gradient as follows: 0 – 1 minute held at 5 % B, 1 – 5 minutes to 8 % B, 5 – 10 minutes to 15 % B, 10 – 13 minutes held at 15 % B, 13 – 23 minutes to 35 % B, 23 – 28 minutes held at 35 % B and back to starting conditions at 29 minutes until 45 minutes. Mass spectrometric parameters were similar for all solvent systems as outlined in section 1.3.1.5 expect for solvent system D under HILIC conditions. The mass spectrometric detection for solvent system D under HILIC conditions at 300 μl/ minute was as follows: The source temperature was set at 350 °C; the spray voltage was 5500V; the declustering potential was set to 50 V for all scans; nitrogen was used as the curtain gas and nebulising gas with flow rates set to 35 AU and 26 AU respectively. Survey scan MS data were acquired by electrospray ionization in positive mode from 400-1200 Da in high resolution mode for 500 ms. Information dependent data acquisition (IDA) was used to collect MS/MS data based on following criteria: the 4 most intense ions with +1 charge and a minimum intensity of 250 cps were chosen for analysis, using dynamic exclusion for 20 seconds after 2 occurrences and a fixed collision energy setting of 47 eV.

For optimisation of two dimensional LCMS work serial coupling of monolith column and HILIC column was set up and solvent A and solvent system D were used as mobile phases. The technique was adapted from the published work used for polar metabolites (Louw, Pereira et al. 2008, Haggarty, Oppermann et al. 2015). For the first separation monolith column (Proswift RP-4H, 1x 250mm) was used using solvent system A as mobile phases and the 2^nd^ separation was performed using HILIC column (ACE Silica, 150 x 3.1mm) column in combination of solvent system D as mobile phases.

The two columns were coupled in series using a T-piece with third port connected to the 2^nd^ binary pump. The scheme of the chromatographic program is explained in appendix 1. The two different gradient program using solvent system A for monolith column and solvent system D for HILIC column are as below:

For reverse phase separation gradient run for solvent system A, the program consisted of a 50 minute run with 0 – 4 minute isocratic hold at 70 % B, followed by 3 step gradient increase to 80 % B at 8 minutes, 90 % B at 15 and 100 % B at 20 minutes. The gradient remained constant until 38 minutes and decreased back to 70 % B until 50 minutes. For HILIC separation gradient consisted of multistep increment as follows: 0 – 1 minute held at 5 % B, 1 – 5 minutes to 8 % B, 5 – 10 minutes to 15 % B, 10 – 13 minutes held at 15 % B, 13 – 23 minutes to 35 % B, 23 – 28 minutes held at 35 % B and back to starting conditions at 29 minutes until 45 minutes.

For the modified gradient program for HILIC column used for 2D-LCMS work the gradient was as follows: 0 – 1 minute held at 5 % B, 1 – 5 minutes to 8 % B, 5 – 10 minutes to 15 % B, 10 – 15 minutes to 17.5 % B, 15 – 20 minutes to 20 % B, 20 – 30 minutes to 30 % B, 30 – 35 minutes to 35 % B, 35 – 40 minutes held at 35 % B and back to starting conditions at 41 minutes until 50 minutes.

## 4 Results & Discussion

The chromatographic separation of OxPC species was evaluated using several columns like C-8 Luna column (150 x 1mm), C-18 Luna column (150 x 1mm), C-30 Luna column (150 x 2.1mm) and Proswift –RP 4H polystyrene – divinyl benzene coated monolith column (1 x 250mm); and mix mode Hichrom amino based HILIC column (150 × 3.1mm) using different solvent systems. Extracted ion chromatogram (XIC) of five representative species of short chain oxidation products, long chain oxidation products and native PCs separated on different columns and solvent system was obtained and the median time point with inter-quartile range of the elution for each group was calculated and graphically represented to evaluate the separation (N=3) (figure 1). The data illustrates that the polystyrene-divinylbenzene coated monolith column provided best separation of short chain oxidation products from long chain oxidation products and unmodified PCs and PEs, compared to other tested columns and solvent systems.

**Figure 1:**
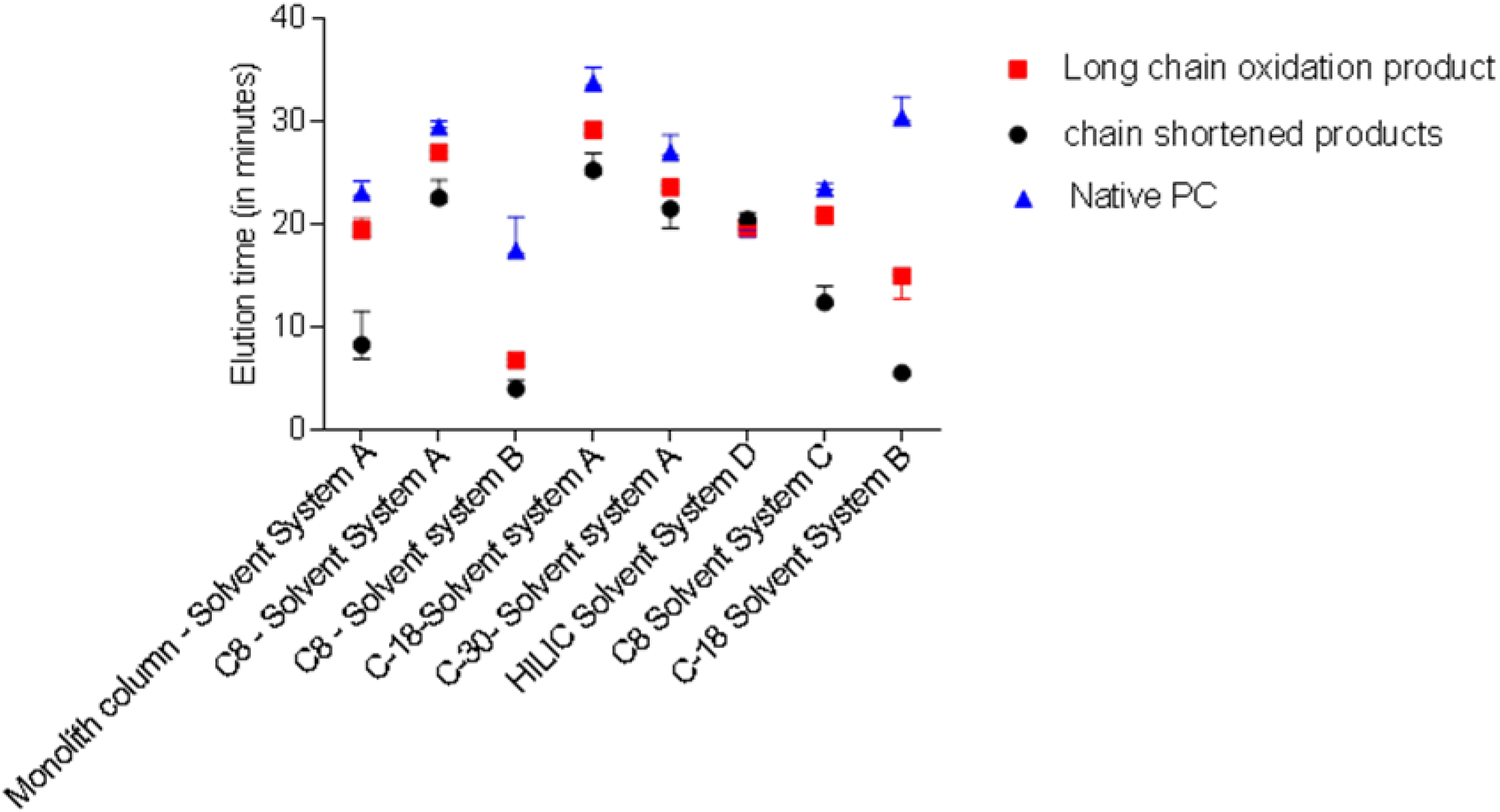
Evaluation of chromatographic separation of OxPC species using several columns and solvent systems. Extracted ion chromatogram (XIC) was generated of five representative species for each group; chain shortened products (POVPC, m/z 594.5; PONPC, m/z 650; SONPC, m/z 678.5; SAzPC, m/z 694.5 and KODA-PC, m/z 704.5), long chain oxidation product (PLPC-OH, m/z 774.5; PLPC-OOH, m/z 790.5; SEIPC, m/z 856.5; SLPC (2OOH), m/z 850.5 and SLPC-OOH, m/z 818.5) and Native PCs (DPPC, m/z 734.5; PLPC, m/z 758.5; SLPC, m/z 786.5; PAPC, m/z 782.5 and SDHPC, m/z 834.5) and the median elution time with inter-quartile range for each group was used to plot the data. Different solvent systems were also used to optimise the separation of different species. **Solvent system B** represents the ternary solvent system consisting of Methanol:Hexane:0.1M ammonium acetate (71:5:7) as eluent A and methanol: hexane (95:5) as eluent B. **Solvent system D** indicates solvent system for HILIC column: eluent A-20% Isopropyl alcohol (IPA) in Acetonitrile (ACN) and eluent B-20 % IPA in ammonium formate (20mM); **Solvent system C** indicates another solvent system consisting of eluent A as Tetrahydrofuran (THF):methanol:water (30:20:50) and eluent B as THF: methanol: water (70:20:10) with 10mM ammonium acetate and **Solvent system A** consist of water with 0.1% formic acid and 5 mM formate as eluent A and methanol with formic acid and 5mM formate as eluent B.

While the C-18 column with Hexane-methanol solvent system (Solvent system B) also provided better separation, reproducibility was an issue owing to volatile nature of the solvent system. Moreover, dispersibility of separation within each group represented by the inter-quartile range, which is defined by the separation of different species belonging to the same group, was observed to be better with monolith column, compared to other conventional reverse phase columns. While the HILIC column separated different phospholipid classes more effectively, different classes of oxidised species within phospholipid class were not separately effectively using a HILIC column. The overlaid extracted ion chromatogram (XIC) of representative species grouped into chain shortened products, long chain oxidation products and native PC’s showing elution profile for different chromatographic separation system is shown.

Serial coupling of the monolith column to HILIC column and running the gradient simultaneously was set up to investigate the improvement in separation of OXPL species. While reverse phase columns separates the polar oxidised species from their native species, it does not separate the oxidised species belonging to different phospholipid class. Moreover, HILIC column provides effective separation of different phospholipid classes but does not effectively separate oxidised species. Moreover, owing to differential ionisation variability, the analysis of different phospholipid classes in a single run is impractical; therefore, with the objective of further improving the separation that can improve the detection range and reduce the analysis time, the evaluation of 2D LC analysis was investigated. The figure 2 shows the separation profile of different groups of oxidised species of PC and PE class when monolith and HILIC column are used for separation alone or in combination by serially coupling the columns and simultaneous gradient analysis. Median elution time of 5 representative species from each group was used to evaluate the separation.

**Figure 2:**
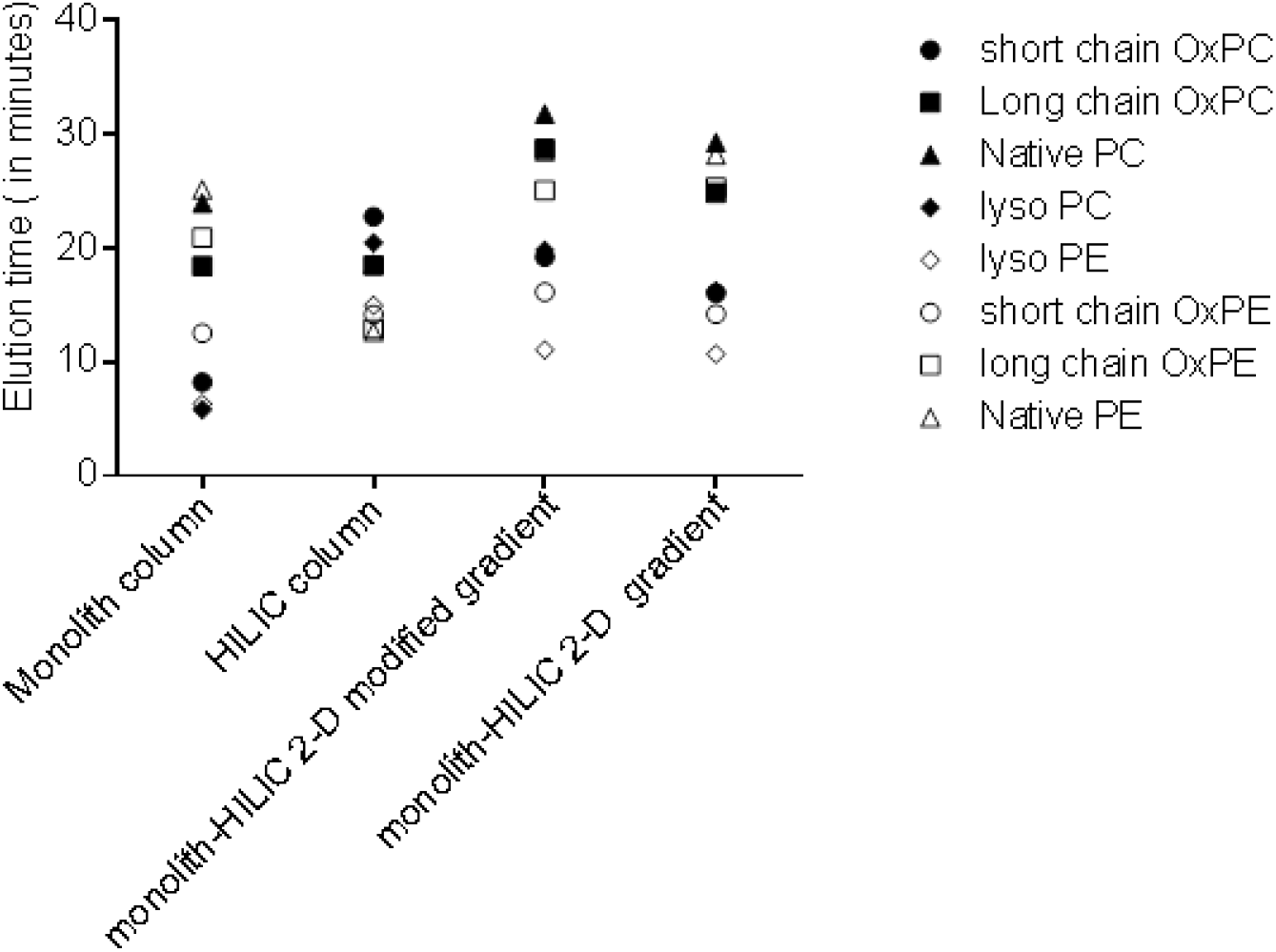
Evaluation of chromatographic separation of OxPC species using monolith and HILIC column alone and in combination by serial coupling and simultaneous gradient analysis. Extracted ion chromatogram (XIC) was generated of five representative species for each group from **PC class:** short chain OxPC (POVPC, m/z 594.5; PONPC, m/z 650.5; SONPC, m/z 678.5; SAzPC, m/z 694.5 and KODA-PC, m/z 704.5), long chain OxPC (PLPC-OH, m/z 774.5; PLPC-OOH, m/z 790.5; SEIPC, m/z 856.5; SLPC (2OOH), m/z 850.5 and SLPC-OOH, m/z 818.5) and Native PCs (DPPC, m/z 734.5; PLPC, m/z 758.5; SLPC, m/z 786.5; PAPC, m/z 782.5 and SDHPC, m/z 834.5); **PE class:** short chain OxPE (POVPE, m/z 552.5; PONPE, m/z 608.5; SONPE, m/z 636.5; SAzPE, m/z 652.5 and KODA-PE, m/z 662.5), long chain OxPE (PLPE-OH, m/z 732.5; PLPE-OOH, m/z 748.5; SEIPE, m/z 814.5; SLPE (2OOH), m/z 808.5 and SLPE-OOH, m/z 776.5) and Native PEs (DPPE, m/z 692.5; PLPE, m/z 716.5; SLPE, m/z 744.5; PAPE, m/z 740.5 and SDHPE, m/z 792.5 and the median elution time for each group was used to plot the data

The serial coupling of monolith and HILIC column with modified program showed better separation of the OxPC and OxPE species. However, the native PCs and PE species co-eluted in this setting. Altogether, this was a first step towards evaluation of two dimensional chromatography to improve the oxidised lipidome coverage and enhance high throughput analysis; and it requires further optimisation and experimentation to evaluate its efficiency.

This was a first attempt to-date in the field of oxidative lipidomics although, similar work had been carried out in the field of metabolomics (Haggarty, Oppermann et al. 2015). We found that the methodology using two dimensional chromatography has a potential to improve the oxidised lipidome coverage as demonstrated by our work and further work needs to be done to evaluate several columns of different chemistry and mobile phases to highlight this not so popular technology in the field of Lipidomics.

